# Asc-Seurat – Analytical single-cell Seurat-based web application

**DOI:** 10.1101/2021.03.19.436196

**Authors:** WJ Pereira, FM Almeida, KM Balmant, DC Rodriguez, PM Triozzi, HW Schmidt, C Dervinis, GJ Pappas, M Kirst

**Affiliations:** School of Forest, Fisheries, and Geomatics Sciences, University of Florida, Gainesville, FL 32611, USA; Department of Cell Biology, Institute of Biological Sciences, University of Brasília, Brasília, DF 70910-900, Brazil; Genetics Institute, University of Florida, Gainesville, FL 32611, USA

## Abstract

**Summary:** Single-cell RNA sequencing (scRNA-seq) has become a popular approach for studying the transcriptome, providing a powerful tool for discovering and characterizing cell types and their developmental trajectories. However, scRNA-seq analysis is complex, requiring a continuous, iterative process to refine the data processing and uncover relevant biological information. We present Asc-Seurat, a feature rich workbench, providing a user-friendly and easy-to-install web application encapsulating the necessary tools for an all-encompassing and fluid scRNA-seq data analysis.

**Availability and implementation:** Asc-Seurat is available at https://github.com/KirstLab/asc_seurat/ and released under GNU 3 license.

**Supplementary information:** Supplementary data are available at *Bioinformatics* online.

## 1. Introduction

Single-cell technologies dramatically enhanced our capacity to characterize tissues and their cell types. By quantifying individual cells’ gene expression, single-cell RNA sequencing (scRNA-seq) substantially increases the resolution of transcriptome profiles from whole tissues or organs by disentangling gene expression signals from bulk cell populations down to cell specific contribution (Stark et al. 2019).

Several software has been developed to address the different aspects of scRNA-seq data analysis. Seurat (Stuart *et al*., 2019) is an R package widely used for scRNA-seq data processing, cell clustering and analysis of differentially expressed genes (DEGs) of single or multiple samples. Further analyses, like single-cell lineage development trajectory inference tools are provided by software such as dynverse (Saelens et al., 2019), in which multiple models can be evaluated and DEGs identified in these trajectories.

The Seurat’s and dynverse’s functions are simple to use, and the results reported are intuitive to interpret. Nevertheless, the dependency on a command-line interface and the need for knowledge of the R programming language poses a major barrier for researchers with limited computational expertise. Moreover, each analytical step outcome is strongly influenced by data quality and execution parameters, requiring the continuous manipulation of these parameters and reevaluation of subsets of data. Tools that simplify this iterative process and integration across analysis platforms are essential for biologists.

Here we present Asc-Seurat (Analytical single-cell Seurat-based web application), an easy to install interactive web application implemented using Shiny (https://shiny.rstudio.com/). Asc-Seurat provides a comprehensive scRNA-seq analysis workbench (Figure 1-A), with the integration of many algorithmic capabilities from Seurat and dynverse, combined with gene functional annotation using BioMart (Smedley *et al*., 2009) via the biomaRt package (Durinck *et al*., 2009).

**Figure 1.**
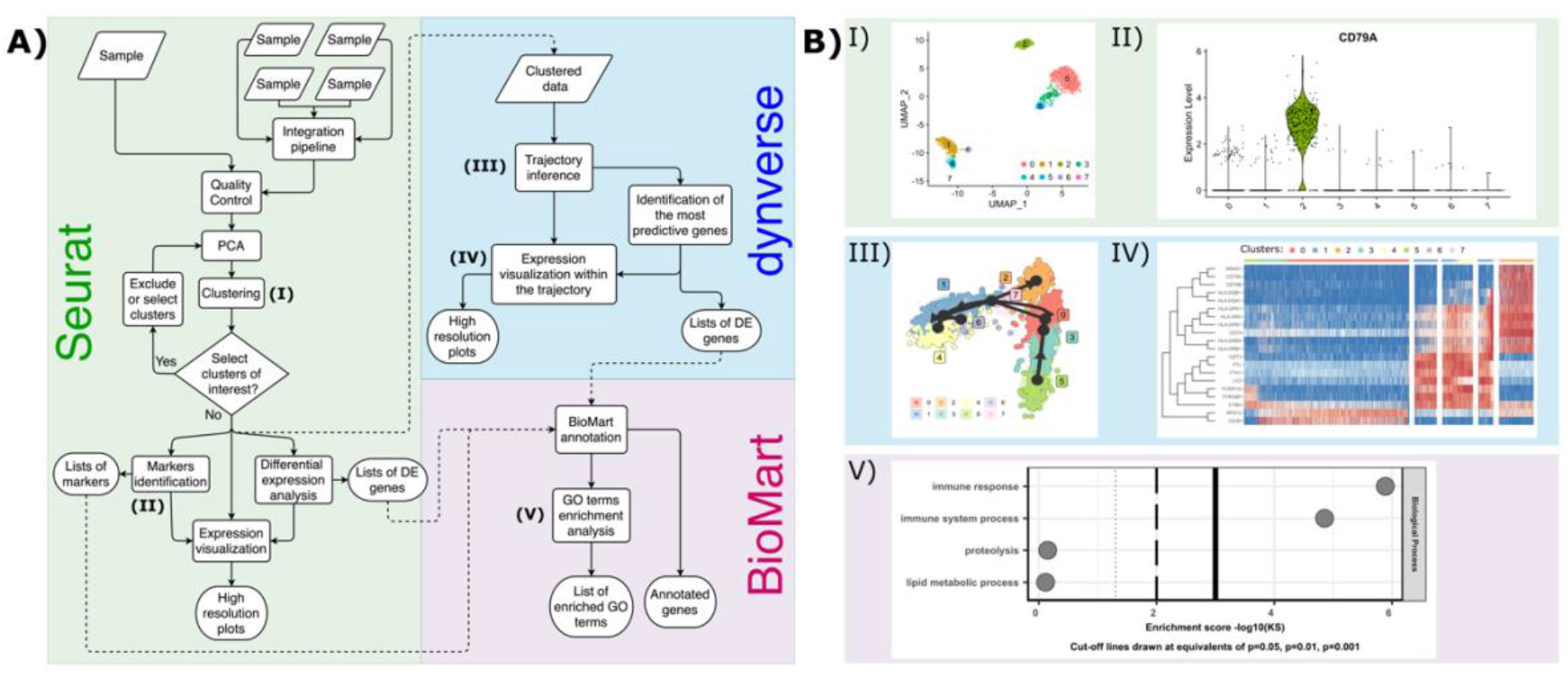
Asc-Seurat workflow overview (A) and demonstrative results using the 10×’s PBMC dataset (B). Asc-Seurat is built on three analytical cores (A). Using Seurat, users explore scRNA-seq data to identify cell types, markers, and DEGs. Dynverse allows the evaluation and visualization of developmental trajectories and identifies DEGs on these trajectories. Asc-Seurat also implements BioMart for functional annotation and GO terms enrichment analysis. B-I) UMAP plot demonstrating the cluster of cells detected in the PBMC sample. B-II) Visualization of the expression of a gene (CD79A) identified as a marker for cluster 2. B-III) Visualization of the developmental trajectory inferred using slingshot (Street *et al*., 2018), a model in dynverse. B-IV) Heatmap showing the gene expression of 20 DEGs, ranked by their importance in predicting the trajectory. B-V) Enriched GO terms, in the category biological process, in the set of 50 most important DEGs within the trajectory shown in B-III.

## 2. Asc-Seurat functionalities

By employing Asc-Seurat, users have access to 1) a rich and responsive graphical user interface, enabling iterative analysis parametrization; 2) a container-based (Docker, https://www.docker.com/) distribution that handles all software dependencies and simplifies installation; 3) integration of multiple samples; 4) selection of cell clusters of interest for reanalysis; 5) search of gene markers and DEG; 6) incorporation of dozens of models for cell trajectory inference; 7) identification of DEGs within trajectories; 8) functional annotation of genes; 9) search for enriched GO terms; and 10) a broad set of publication-ready graphs, among other features.

## 3. Asc-Seurat use case

To demonstrate Asc-Seurat’s functionalities, we analyzed the publicly available 10×’s Peripheral Blood Mononuclear Cells (PBMC) dataset (**Figure 1-B)**. A stepwise demonstration can be found in the **Supplementary material**.

### 3.1. Quality control, clustering, and markers identification

Asc-Seurat workflow starts with raw scRNA-seq data input and provides several options to exclude poor quality cells. Violin plots are generated to show the distribution of cells before and after filtering (**Figure S2**). Expression levels can be normalized and scaled (Figure S3), followed by clustering and cluster visualization using UMAP and t-SNE (**Figure 1.B-I and Figure S4**). After clustering, Asc-Seurat simplifies the selection and reanalysis of subsets of cells from clusters of interest. Moreover, the expression of genes can be visualized at the individual cell (**Figure S7**) or cluster level (**Figure 1.B-II and Figure S8**). Asc-Seurat also implements methods to identify cell-type markers and genes differentially expressed among clusters (**Figure S5**).

### 3.2. Trajectory inference and differential expression

Several algorithms are available on Asc-Seurat to model and infer developmental trajectories among cell clusters (**Figure 1.B-III and Figure S11**), allowing the visualization of gene expression in individual cells within the trajectory (**Figure S13**) and the identification of DEGs along the trajectory **(Figure 1.B-IV, Figure S12)**.

### 3.3. Functional annotation and enrichment analysis

Annotation of genes of interest (e.g., markers, DEGs) can be retrieved by the Asc-Seurat BioMart module. Moreover, it is possible to search for enriched Gene Ontology (GO) terms (**Figure 1.B-V and Figure S15)** using topGO (Alexa and Rahnenfuhrer, 2020; Bioconductor). These capabilities allow the biological interpretation of scRNA-seq results and real-time hypotheses generation.

## Data availability

The 10×’s PBMC dataset is publicly available at https://cf.10xgenomics.com/samples/cell/pbmc3k/pbmc3k_filtered_gene_bc_matrices.tar.gz

## References

Alexa A, Rahnenfuhrer J (2020). topGO: Enrichment Analysis for Gene Ontology. R package version 2.42.0.

Durinck, S. et al. (2009) Mapping identifiers for the integration of genomic datasets with the R/Bioconductor package biomaRt. Nat. Protoc., 4, 1184–1191.

Saelens, W. et al. (2019) A comparison of single-cell trajectory inference methods. Nat. Biotechnol., 37, 547–554.

Smedley, D. et al. (2009) BioMart--biological queries made easy. BMC Genomics, 10, 22.

Street, K. et al. (2018) Slingshot: cell lineage and pseudotime inference for single-cell transcriptomics. BMC Genomics, 19, 477.

Stark, R. et al. (2019) RNA sequencing: the teenage years. Nat. Rev. Genet., 20, 631–656.

Stuart, T. et al. (2019) Comprehensive Integration of Single-Cell Data. Cell, 177, 1888–1902.e21.

