## Supplementary material for "Asc-Seurat – Analytical single-cell Seurat-based web application"

#### Supplementary Information

Pereira WJ[1], Almeida FM[2], Balmant KM[1], Rodriguez DC[1], Triozzi PM[1], Schmidt HW[1], Dervinis C[1], Pappas Jr GJ[2], Kirst M[1,3*]

[1] School of Forest, Fisheries, and Geomatics Sciences, University of Florida, Gainesville, FL 32611, USA.

[2] Department of Cell Biology, Institute of Biological Sciences, University of Brasília, Brasília, DF 70910-900, Brazil.

[3] Genetics Institute, University of Florida, Gainesville, FL 32611, USA.

*To whom correspondence should be addressed.

As a case study, we analyzed the 10×'s Peripheral Blood Mononuclear Cells (PBMC) 3k dataset.

#### Introduction

Asc-Seurat provides a rich and responsive single-cell RNA-seq (scRNA-seq) analysis workbench, designed to facilitate the overall execution of multiple steps of a typical scRNA-seq study. This document provides a summary of Asc-Seurat's capabilities and a showcase of its user interface and graphical output illustrating the selection of analysis parameters and interpretation of results. Below, for the sake of simplicity, we demonstrate the main steps of analysis for an individual sample. However, a multiple sample option is also available on Asc-Seurat by deploying Seurat’s integration algorithm.

A full instruction manual guiding the installation and a more comprehensive description of Asc-Seurat capabilities, such as parameter options, available algorithms, and user interface is provided at readthedocs.io.

### 1. Analysis of the Peripheral Blood Mononuclear Cells dataset

To generate the results presented below, we analyzed the 10×'s PBMC 3k dataset, available at https://cf.10xgenomics.com/samples/cell/pbmc3k/pbmc3k_filtered_gene_bc_matrices.tar.gz.

#### 1.1. Loading the data

For analysis using Asc-Seurat, all scRNA-seq datasets should be stored in a folder inside the folder “*data/*”, which is generated during the installation. Asc-Seurat will display compatible files stored within the “*data/*” folder, from where the data of interest can be selected. Next, the user can provide a name for the project and define the initial parameters to select cells to be loaded in the web application (**Figure S1**).

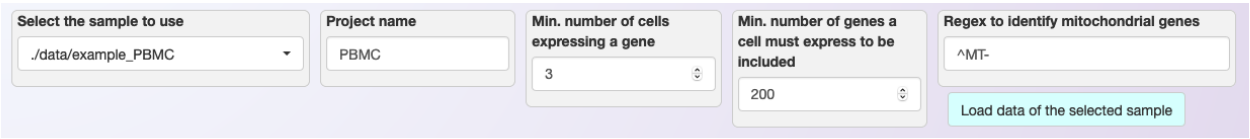


**Figure S1. Loading the dataset on Asc-Seurat.** From the stored datasets, users can select the sample to analyze and define the minimal cell requirements to load the data. For the 10×'s PBMC, only cells expressing at least 200 genes and only genes expressed in three or more cells were considered.

#### 1.2. Quality Control

After loading the dataset, a violin plot shows the distribution of the number of expressed genes (nFeature_RNA), the number of Unique Molecular Identifiers or independent transcript (nCount_RNA), and the percentage of mitochondrial genes (percent.mt) detected in each cell. Users can define more restrictive parameters to remove undesirable cells. A second violin plot showing the filtered dataset allows users to visualize the filtering effect and adjust the parameters accordingly (**Figure S2**).

As for all plots generated by Asc-Seurat, it is possible to download a high-resolution image containing the violin plot displaying the filtered dataset.

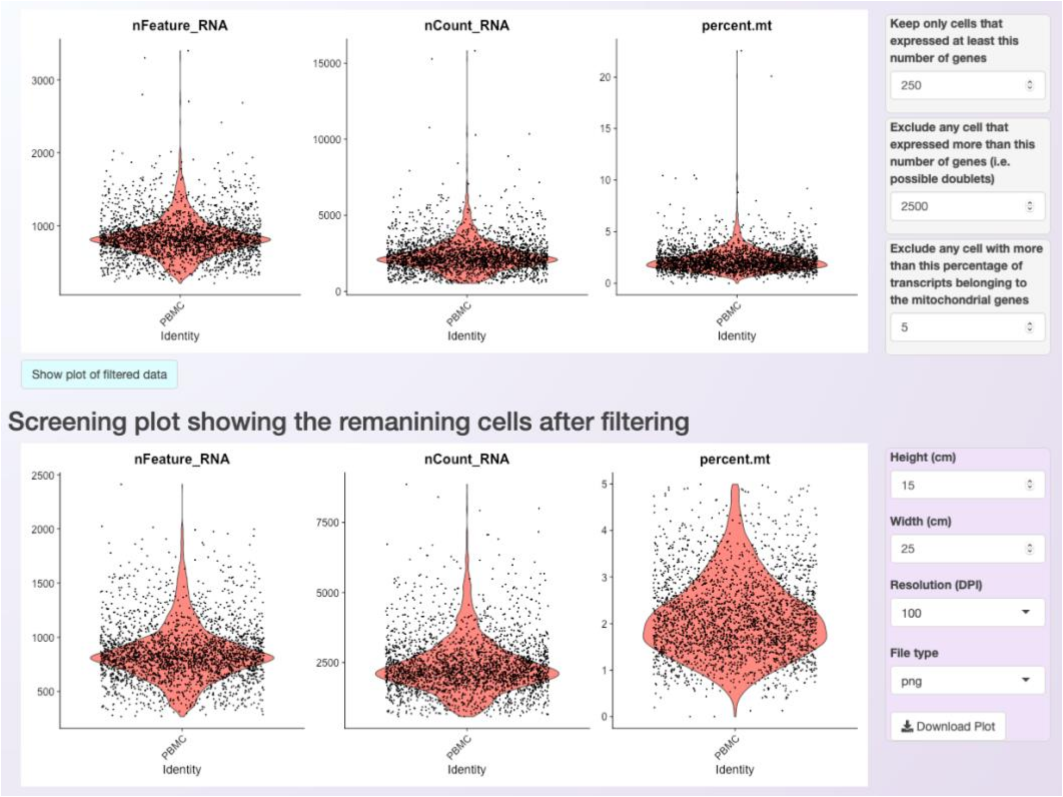


**Figure S2. Data filtering.** Violin plots showing the distribution of cells before (upper panel) and after (lower panel) the filtering of undesirable cells.

##### 1.3. Scaling, normalization, and dimension reduction using Principal Component Analysis (PCA)

Next, users can select the scale factor for Seurat’s log normalization. Also, it is possible to define how many of the most variable genes should be used in the posterior steps, and what method should be used to identify the most variable genes (**Figure S3**).

After a user starts the PCA analysis by clicking on “Run the PCA analysis”, the dataset will be normalized, scaled and the PCA analysis will be generated using the selected number of most variable genes. An elbow plot showing the contribution of the principal components (PC) to the variation in the dimensionally reduced space is generated (**Figure S3**) to help users decide how many PCs should be used during the clustering step.

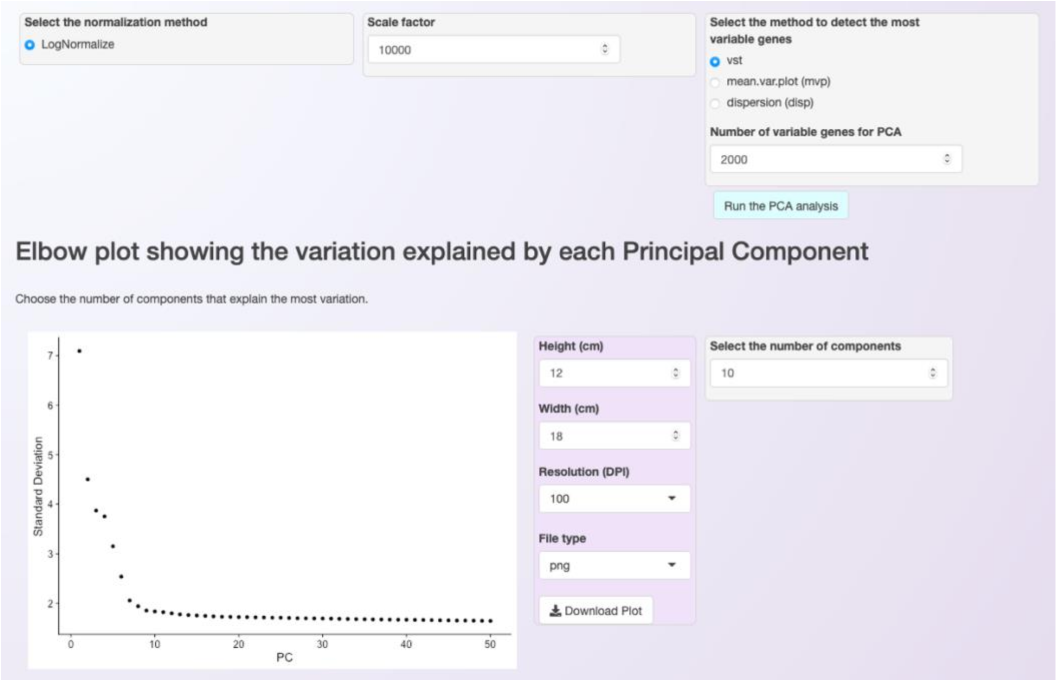


**Figure S3. Asc-Seurat integrates the normalization, scaling, and PCA in one step.** After choosing the required parameters (top), users can trigger the analysis and Asc-Seurat will execute the normalization, identification of the most variable genes, and PCA. An elbow plot (bottom left) is then generated to help users select the number of PCs to be used in the clustering. For the PBMC dataset, the first 10 PCs (bottom right) were selected.

#### 1.4. Clustering

After choosing the PCs to be used, it is necessary to inform what resolution should be used during the clustering. The resolution is an important parameter to evaluate because it determines the profile and number of clusters identified for a dataset. Selecting larger values will favor splitting cells into more clusters while selecting a smaller value has the opposite effect.

For the PBMC dataset, a resolution of 0.5 was selected and nine clusters were identified (**Figure S4**).

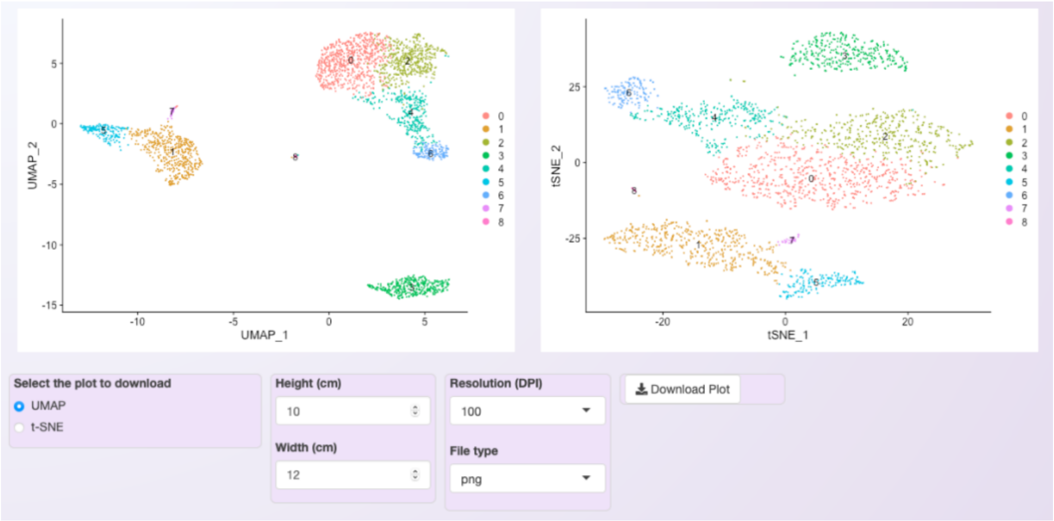


**Figure S4. Two plots are available for cluster visualization in Asc-Seurat.** The first plot is generated using the Uniform Manifold Approximation and Projection (UMAP) technique (left).

The second deploys the t-distributed Stochastic Neighbor Embedding (t-SNE) method (right). Nine clusters were obtained for the PBMC dataset.

##### 1.5. Differential Expression Analysis

Asc-Seurat provides an assortment of algorithms to identify gene markers for individual clusters or to identify differentially expressed genes (DEGs) among clusters. For the PBMC dataset, when using the Wilcox test, filtering for genes that are expressed in at least 10% of the cells in the cluster, with a (log) fold change higher than 0.25 and an adjusted p-value smaller than 0.05, almost 400 gene markers were identified for cluster 3 (Figure S5).

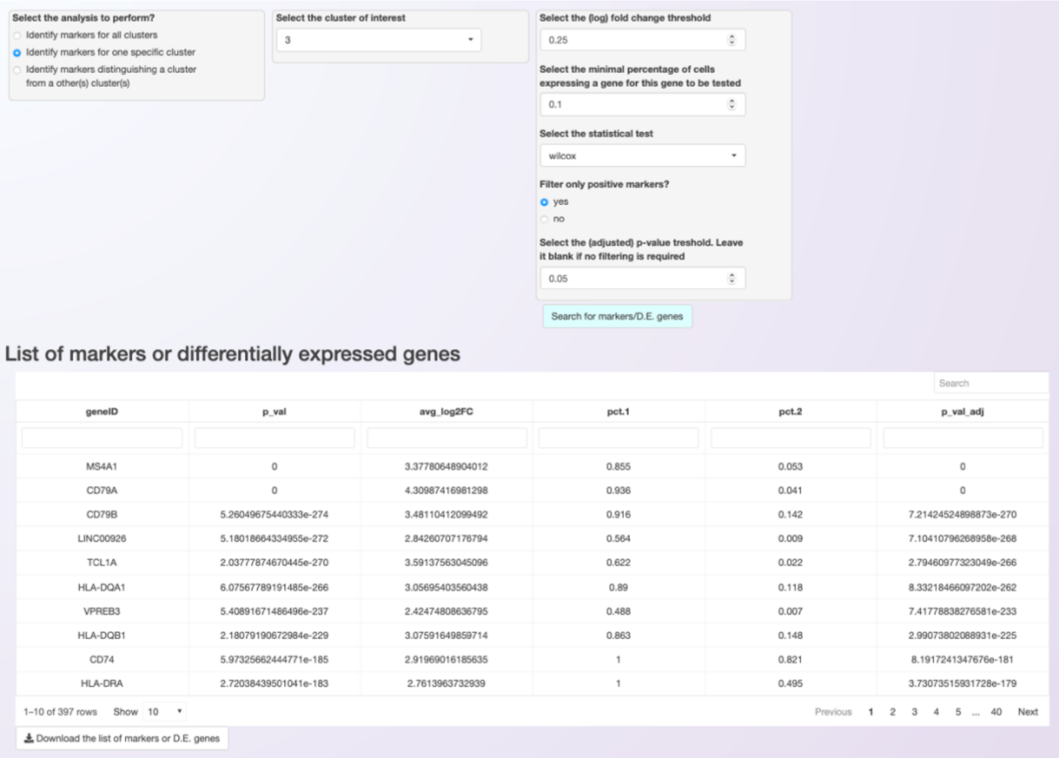


**Figure S5. Asc-Seurat allows the identification of gene markers and DEGs**. Users can search for gene markers for all clusters, for a specific cluster, or search for DEGs among specific clusters (top). An iterative table displaying the significant genes is generated (bottom), and users can download the list of markers, or DEGs, as a csv (comma-separated values) file.

The list of significant genes can be download and the resulting csv file used as input for the sessions of the web application dedicated to the visualization of gene expression, or to the functional annotation using BioMart (see below).

#### 1.6. Gene expression visualization

Asc-Seurat provides a variety of plots for gene expression visualization. From a list of selected genes, it is possible to visualize in a heatmap the average of each gene expression in each cluster (**Figure S6**) and at the cell level (**Figure S7**). Moreover, violin plots and dot plots allow the visualization of each cluster's expression, with emphasis on inter-cluster comparison (**Figure S8**).

##### 1.6.1. Heatmap

Using the list of markers identified for PBMC's cluster 3 as input, we selected the five most DEGs for visualization in the heatmap (**Figure S6**).

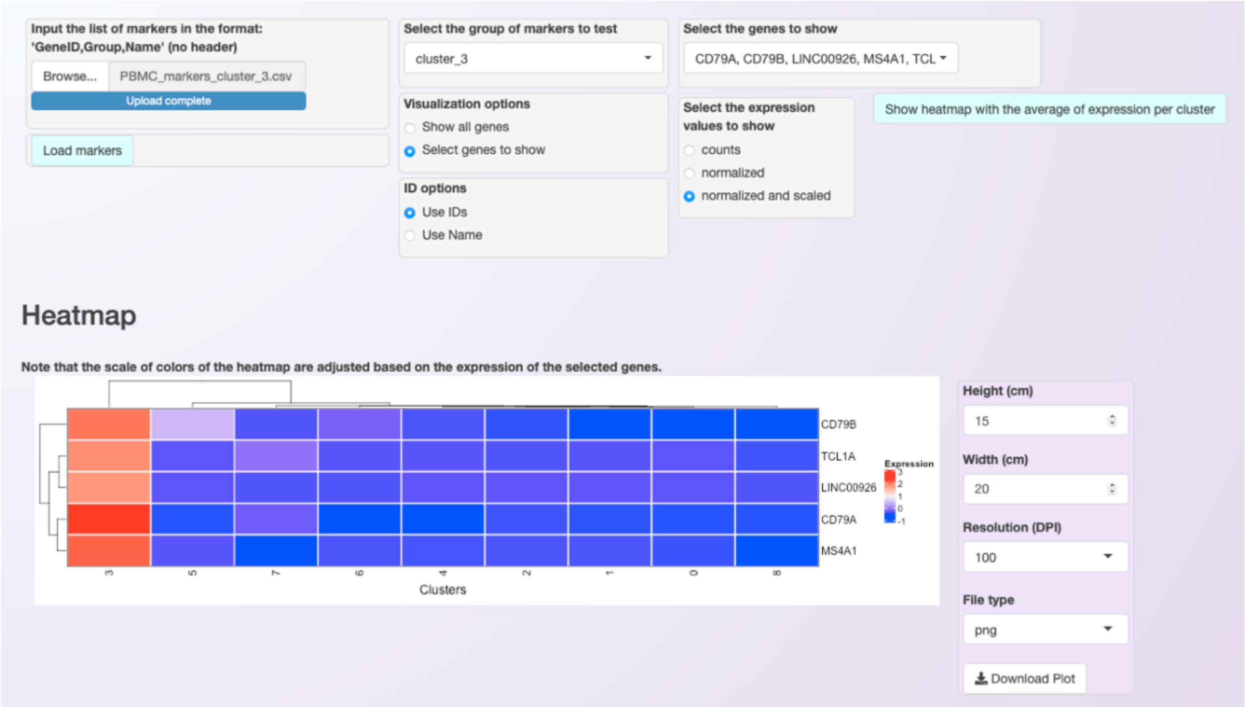


**Figure S6. Asc-Seurat provides a heatmap to allow the comparison of the gene expression profile among clusters.** Users can provide a list of genes as a csv file, filter those by group, and/or select the genes of interest (top). The heatmap shows the average expression of the selected genes in each cluster. Moreover, Asc-Seurat groups genes with similar expression using a hierarchical clustering algorithm. Asc-Seurat adjusts the plot height based on the number of selected genes, and it can be readily downloaded (bottom).

For the five selected genes, the expression profile is clearly pronounced on cluster 3, confirming that they are useful markers for this cluster.

##### 1.6.2. Expression visualization per cell

After visualizing the expression profiles in the heatmap, the user can select genes to explore in more detail. For each selected gene, Asc-Seurat generates plots that allow the visualization of the gene expression at the cell level (**Figure S7)**.

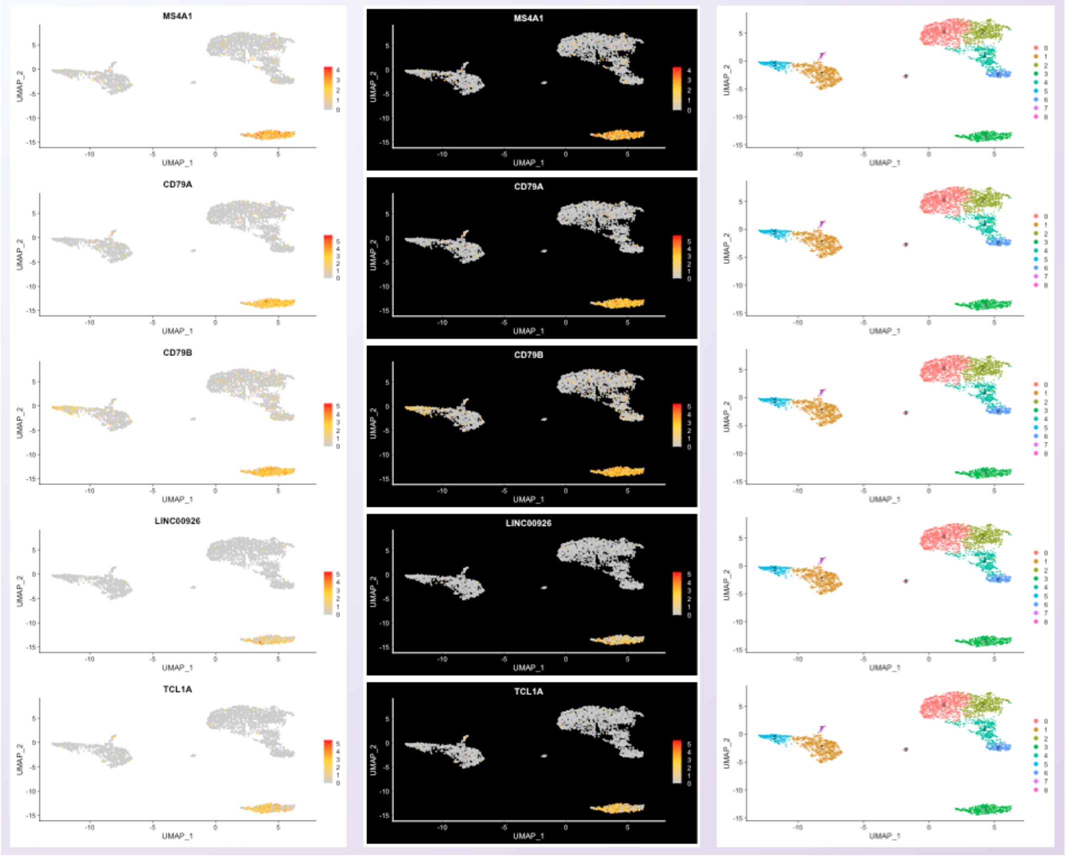


**Figure S7. Asc-Seurat allows the visualization of the gene expression at the cell level by deploying Seurat's feature plots.** Using the same five markers selected for cluster 3 on the heatmap, it is possible to observe that the expression of these genes is, as expected, higher among cells in the cluster 3, with a large percentage of cells of this cluster expressing the selected genes.

##### 1.6.3. Visualization of the expression among clusters

Asc-Seurat also allows the visualization of the distribution of cells within each cluster according to the expression of the gene (violin plot) and the percentage of cells in each cluster expressing the gene (dot plot) (**Figure S8**).

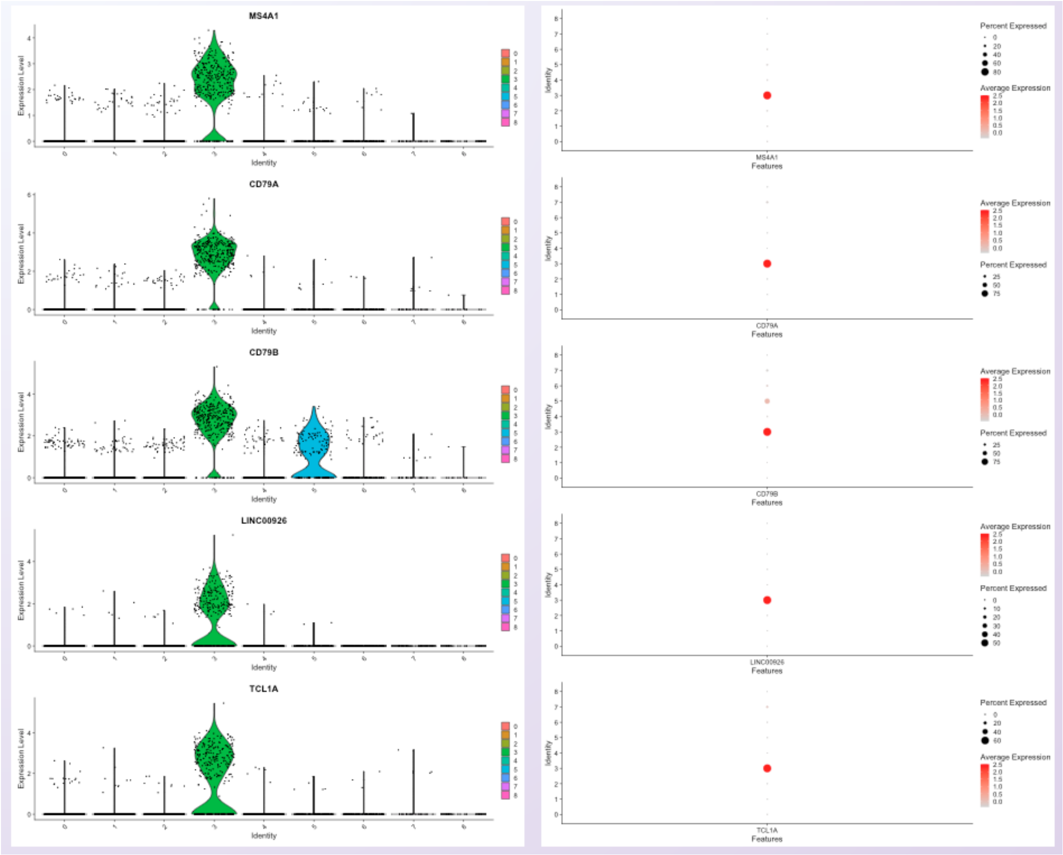


**Figure S8. Asc-Seurat allows an easy way to compare the expression profile of a gene among clusters.** Violin plots show the distribution of cells within each cluster according to the gene expression (left panel), and dot plots show the percentage of cells in each cluster expressing the gene (right panel).

Using the same five gene markers identified for cluster 3, violin and dot plots confirm that their expression is detected in the expected cluster. However, by deploying the violin and dot plots, it is possible to observe that one of the markers (CD79B) is also significantly expressed on cluster 5, which was not distinguishable in the heatmap.

#### 1.7. Trajectory inference

For trajectory inference analysis, users can either execute it through capabilities of the embedded slingshot R package  or select another model contained in dynverse, executed using a docker image provided by the later. In both options, users only need to select the model and initial parameters (see below). However, the direct execution of slingshot is faster than the execution of models via dynverse's docker image.

##### 1.7.1. Trajectory visualization

To start the trajectory inference analysis, users need to save the clustered data in a specific folder automatically created during the installation. Asc-Seurat recognizes the data automatically and users can select the sample to be used. Next, users need to select the model to be used, inform if the data is composed by one or multiple integrated samples and, optionally, inform the cluster(s) expected to be at the beginning and/or end of the inferred trajectory. After executing the analysis, three plots showing different inferred trajectory representations are generated (**Figure S11**).

For the PBMC dataset, a developmental trajectory containing three lineages was identified.

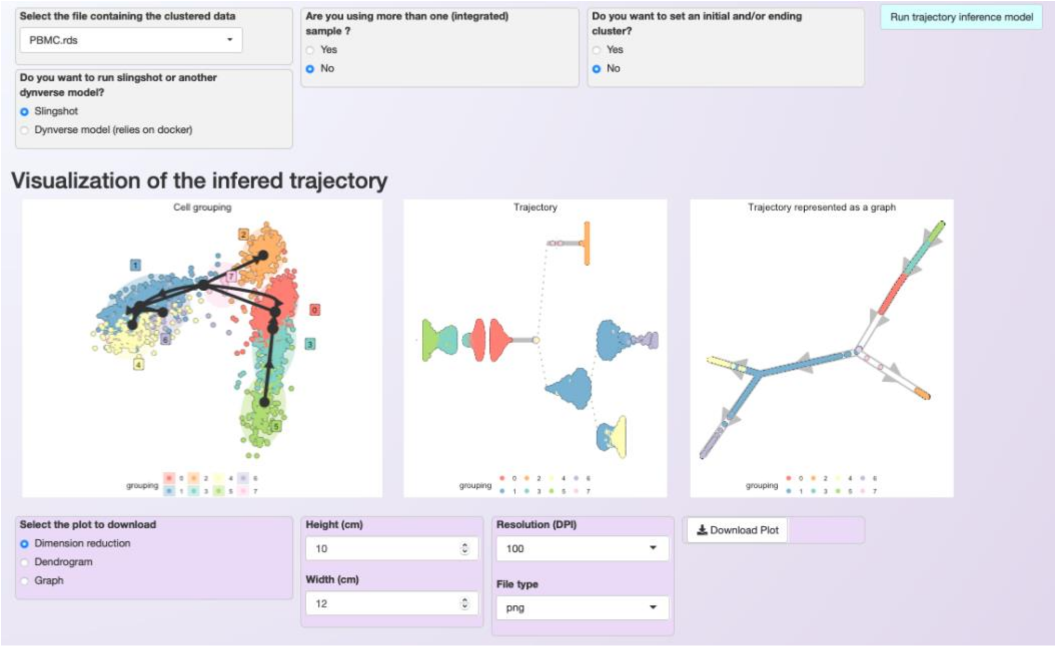


**Figure S11. Asc-Seurat provides multiple models for trajectory inference analysis and three options for trajectory visualization.** Users only need to choose the dataset, the model and, optionally, to inform the cluster(s) that are expected to be at the start and/or at the end of the trajectory (top). Three representations of the trajectory are shown, facilitating the interpretation of results (bottom). Moreover, if the dataset corresponds to the integration of multiple samples, it is possible to change the color scheme so cells are colored by sample instead of by cluster.

##### 1.7.2. Gene expression within the trajectory

After inferring the developmental trajectory, it is possible to visualize the expression of genes of interest in the cells within the trajectory. Asc-Seurat provides two options for this visualization, 1) a heatmap displaying the expression of genes in each cell, ordered by the cell position within the trajectory (**Figure S12**), and 2) the visualization of the same three trajectory's representation shown above but colored by the gene expression (**Figure S13**).

User can either load their list of genes of interest or identify DEGs within the trajectory for the visualization. For the PBMC dataset, we opted to show the 50 most significant DEGs within the trajectory, as ranked by their "importance" value on explaining the inferred trajectory (**Figure S12**).

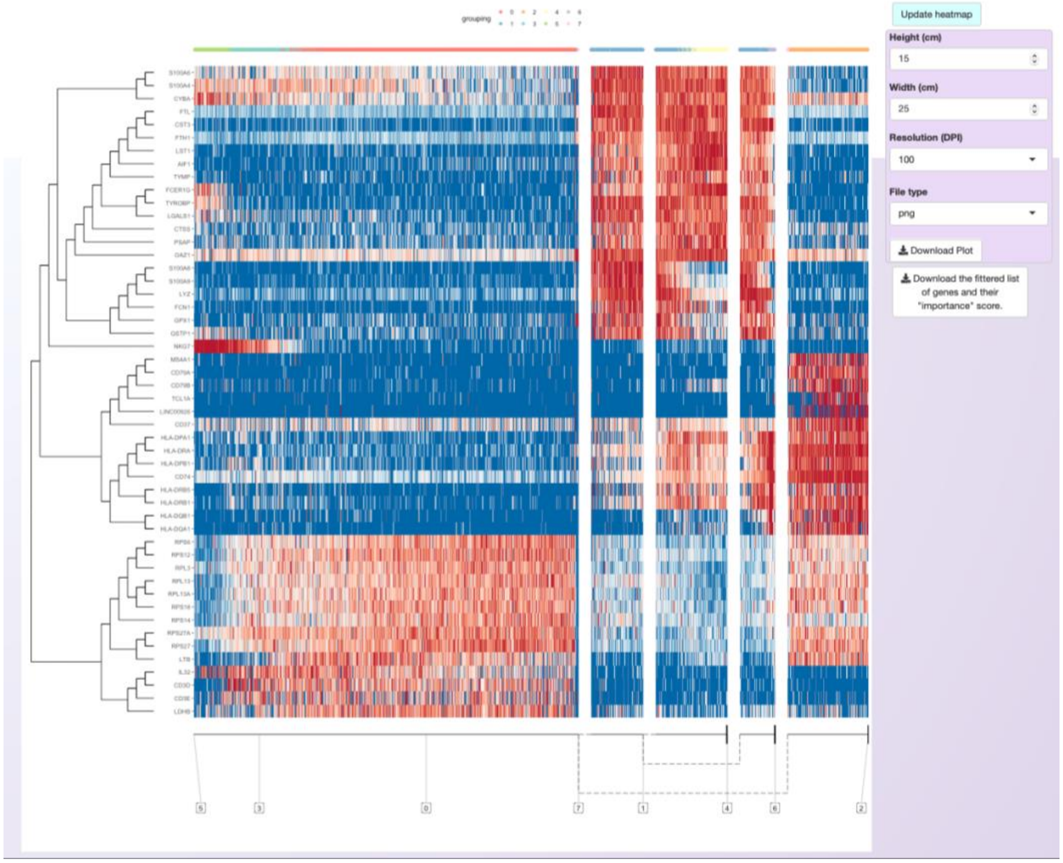


**Figure S12. A heatmap showing the expression of genes within cells by the order that these cells appear in the trajectory.** At the bottom of the heatmap, a representation of the branches of the trajectory facilitates the interpretation of the gene expression profiles. In the top of the heatmap, clusters are represented by the same color scheme as in the trajectory representation described previously.

From the genes included in the heatmap, users can select a subset to visualize their expression using the three trajectory representation described above.

For the PBMC dataset, we selected three genes as examples. The gene NKG7 is expressed in cells in the beginning of the trajectory, while transcripts for MS4A1 and are detected more specifically in alterantive branches in later parts of the trajectory (**Figure S13)**.

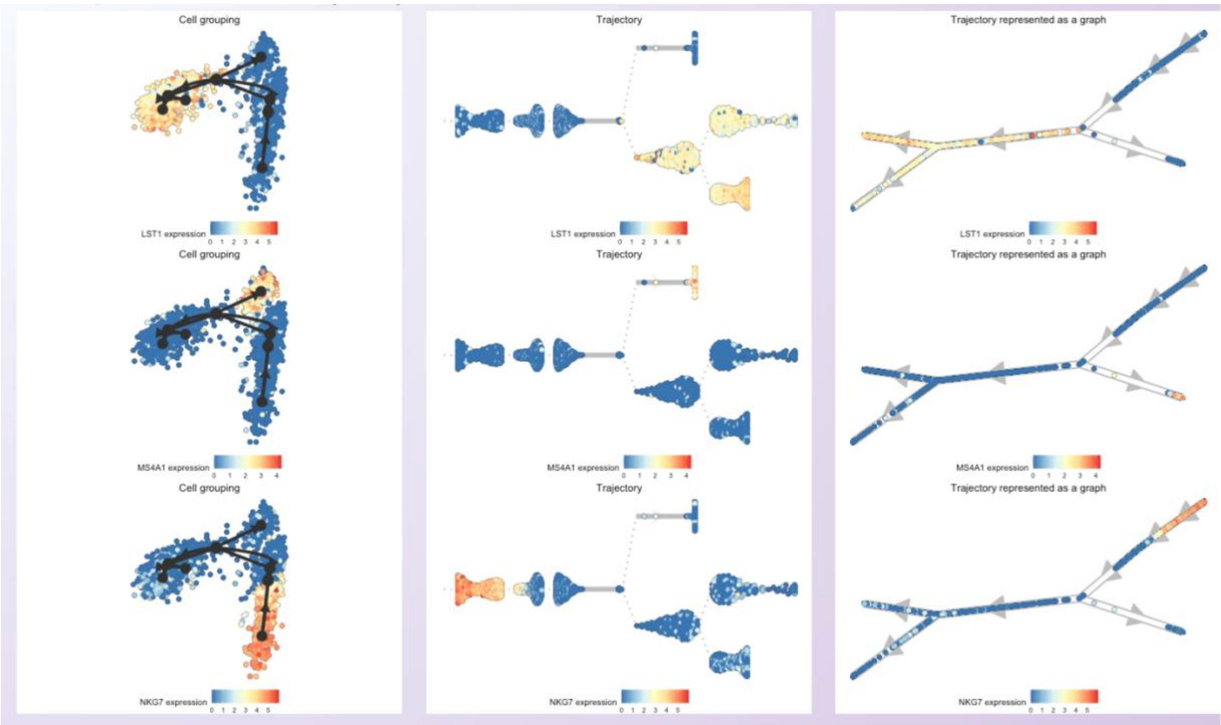


**Figure S13. Asc-Seurat allows users to select genes to visualize their expression profile at the cell level and within the trajectory.** For each gene, three representation of the trajectory are generated showing the expression of the genes in the cells ordered in the trajectory.

#### 1.8. Functional annotation and Gene Ontology enrichment analysis

In many instances, users are interested in obtaining more information about a gene, or a set of genes, to support the interpretation of the data and development of new hypotheses. For example, Asc-Seurat produces lists of gene markers, DEGs or DEGs within a trajectory that might be of special interest. Next, functional annotation for these genes can be retrieved by querying BioMart servers (for several species) and, finally, Gene Ontology (GO) term enrichment analysis to verify if one or more GO terms are over-represented or under-represented in a set of selected genes.

For example, the GO terms enrichment analysis of the 50 most important DEGs within the trajectory inferred for the PBMC dataset identifies two terms related to the immune system as the most significantly enriched terms (**Figure S15**).

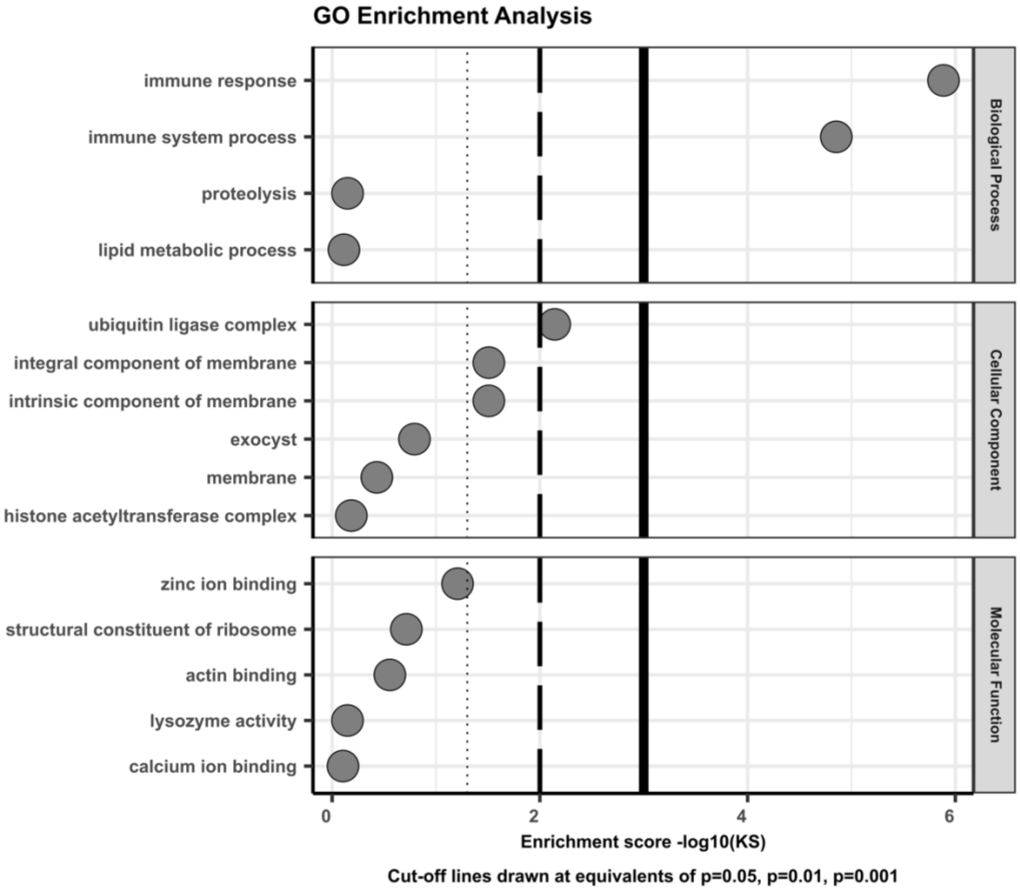


**Figure S15. Asc-Seurat provides the functional annotation of genes from a myriad of species via BioMart. It also provides the Gene Ontology term enrichment analysis via topGO, a Bioconductor package.** Users can download the list of enriched GO terms and/or generate the plot above for the most significant enriched GO terms.
